# Rectal swabs in critically-ill patients provide discordant representations of the gut microbiome compared to stool samples: a brief methodologic report

**DOI:** 10.1101/541441

**Authors:** Katherine Fair, Daniel G. Dunlap, Adam Fitch, Tatiana Bogdanovich, Barbara Methé, Alison Morris, Bryan J. McVerry, Georgios D. Kitsios

## Abstract

The role of the gut microbiome in critical illness is being actively investigated, but the optimal sampling methods for sequencing studies of gut microbiota remain unknown. Stool samples are generally considered reference-standard but are not practical to obtain in the intensive care unit (ICU), and thus, rectal swabs are often used. However, the reliability of rectal swabs for gut microbiome profiling has not been established in the ICU setting. In this study, we compared 16S rRNA gene sequencing results between rectal swab and stool samples collected at three timepoints in mechanically-ventilated critically-ill adults. Rectal swabs comprised 89% of samples collected at the baseline timepoint, but stool samples became more available at later time-points. Significant differences in alpha and beta-diversity between rectal swabs and stool samples were observed, but these differences were primarily due to baseline samples. Higher relative abundance of Actinobacteria phyla (typically skin microbes) was present in rectal swabs compared to stool samples (p=0.05), a difference that was attenuated over time. The progressive similarity of rectal swabs and stool samples likely results from increasing stool coating of the rectal vault and direct soiling of the rectal swabs taken at later time points. Therefore, inferences about the role of the gut microbiome in critical illness should be drawn cautiously and take into account the actual type and timing of samples analyzed.

**Statement of Importance:** Rectal swabs have been proposed as potential alternatives to stool samples for gut microbiome profiling in outpatients or healthy adults, but their reliability in critically-ill patients has not been defined. Because stool sampling is not practical and often not feasible in the intensive care unit, we performed a detailed comparison of gut microbial sequencing profiles between rectal swabs and stool samples in a longitudinal cohort of critically-ill patients. We identified systematic differences in gut microbial profiles between rectal swabs and stool samples and demonstrated that timing of rectal swab sampling had a significant impact on sequencing results. Our methodological findings should provide valuable information for the design and interpretation of future investigations of the role of the gut microbiome in critical illness.

## Introduction

Gut microbial dysbiosis is a plausible contributor to the onset, evolution and outcome of critical illness, but the mechanisms involved have not been fully elucidated (1,2). Fecal microbial communities in critically-ill adults display lower diversity and distinct taxonomic signatures compared to fecal samples from healthy individuals (3). Thus, defining the pathogenetic disruptions of gut communities during critical illness may help identify new targets for intervention (1,2).

Sampling gut microbiota for sequencing analyses can be challenging in the intensive care unit (ICU). Critically-ill patients frequently experience constipation or ileus (4), and the provision of early enteral nutrition is highly variable (5). Thus, critically-ill patients may not have any bowel movements, especially early in their ICU course. Furthermore, standard decontamination practices in ICU care (6) often result in stool disposal before samples are collected. For these reasons, rectal swabs represent an attractive, minimally invasive method for sampling gut microbiota, which is routinely used in clinical practice for Vancomycin-Resistant *Enterococcus* colonization screening.

Rectal swabs have been proposed as potential alternatives to stool samples in ambulatory patients (7,8), but data in critical illness are limited. Recent work from Bansal et al. (9) in nine critically-ill patients showed compositional discrepancies between rectal swabs and stool samples when rectal swabs were not visibly soiled by stool. To further examine for systematic differences in gut microbial profiles captured by stool vs. rectal swab samples, we obtained data from a larger cohort of 106 patients admitted to the medical ICU at a tertiary academic center.

## Methods

Detailed methods are reported in the Supplement. Briefly, in this observational cohort study, we prospectively enrolled consecutive mechanically-ventilated patients with acute respiratory failure from any cause. We collected rectal swabs and/or stool samples at baseline [days 0-2 form intubation], middle [days 3-6] and late [days 7-10] intervals of follow-up starting at the time of intubation and continuing for up to 10 days if the patient remained in the ICU. Rectal swabs were collected according to a standard operating procedure (i.e. placing the patient in a lateral position, entering the cotton tip of the swab in the rectal canal and rotating gently for five seconds), unless clinical reasons precluded movement of the patient (e.g. severe hemodynamic or respiratory instability). Stool samples were collected when available, either by taking a small sample from an expelled bowel movement (before cleaning of the patient and disposal of stool), or through a fecal management system (rectal tube) placed for management of diarrhea and liquid stool collection. For comparisons with healthy gut microbiota, we also included 15 stool samples obtained from healthy volunteers used for fecal microbiota transplantation (FMT stool). We extracted bacterial DNA and performed 16S rRNA gene sequencing (V4 subunit) on Illumina MiSeq with standard protocols as previously described and detailed in the Supplement (10). Sequencing data were analyzed for alpha/beta-diversity and taxonomic composition with the R software.

## Results

We enrolled 106 patients with a total of 171 samples (132 rectal swabs and 39 stool samples). Stool samples were available from 25 (24%) patients during the study period, and 10 patients had both sample types available. Patients with stool samples available had similar baseline demographics (age, sex, BMI) compared to patients without stool samples (Figure 1A), but had higher severity of illness by Sequential Organ Failure Assessment (SOFA) scores and longer duration of ICU stay and mechanical ventilation (Wilcoxon p<0.001). Sample type availability varied by follow-up interval: rectal swabs constituted the majority of samples at the baseline interval (87%), but stool samples were progressively available for larger proportions of patients (61% of patients at late interval, Figure 1B).

**Figure 1:**
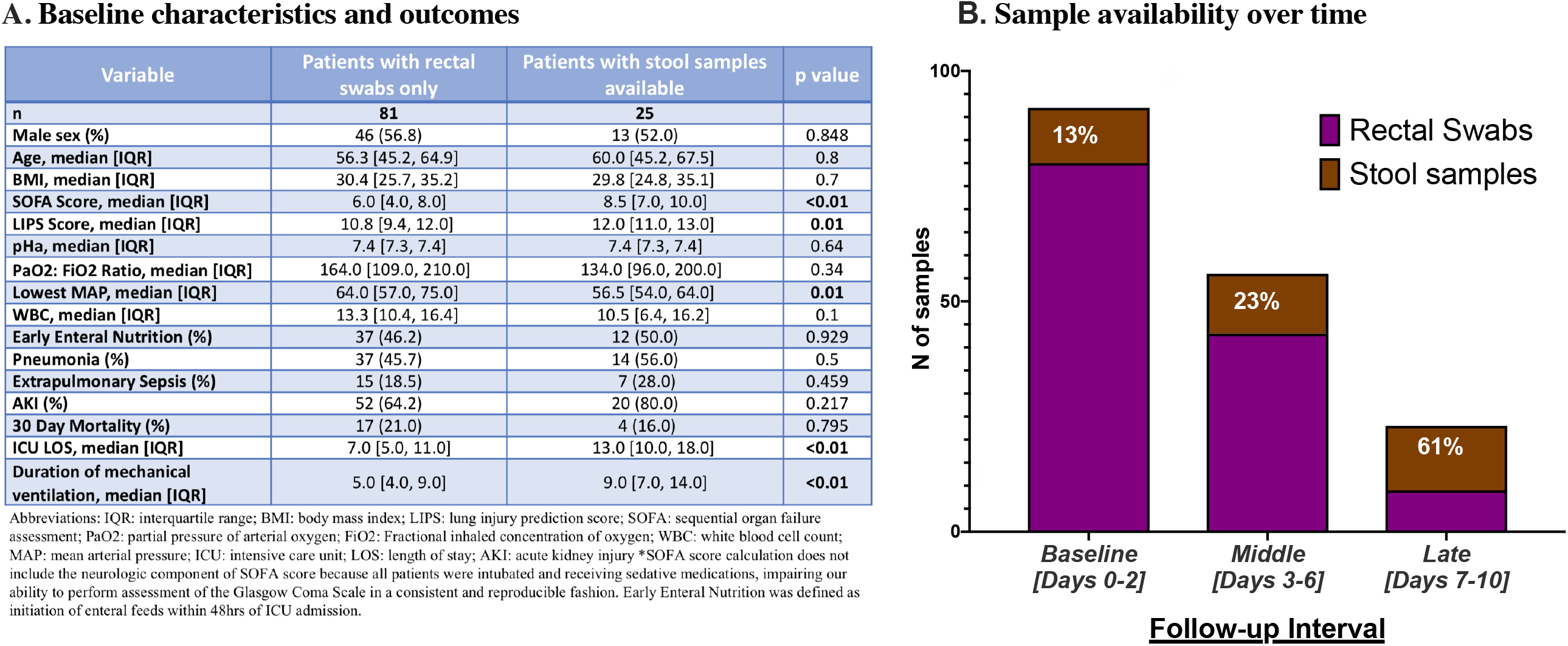
Cohort characteristics and sample type availability over time. A. Table with baseline characteristics and clinical outcomes of patients with rectal swabs only vs. patients with stool samples available. P-values are from Wilcoxon tests for continuous and Fisher’s exact tests for categorical variables (highlighted in bold when significant p<0.05). B. Stacked bar-graph of number of rectal swabs vs. stool samples at each time interval (purple for rectal swabs and brown for stool samples). The proportion of stool samples available at each time interval is shown in white.

Rectal swabs and stool samples overall produced similar number of reads (high quality 16S rRNA gene sequences, median [interquartile range-IQR]= 4235[1034]), which was much higher than the number of reads produced by experimental negative controls (Wilcoxon p<0.0001, Figure S1), suggesting successful recovery of bacterial DNA signal by both sampling techniques. In the baseline interval, rectal swabs had higher alpha-diversity (Shannon= 2.4[1.7]) compared to stool samples (3.1[1.3], Wilcoxon p=0.02) (Figure 2A and multivariate adjusted analysis in Table S1), but samples collected later had similar alpha-diversity (p=0.68 and 0.30 for middle and late interval comparisons, respectively). Notably, both rectal swabs and stool samples had significantly lower alpha-diversity compared to FMT stool samples from healthy donors (Wilcoxon p<0.0001, Figure 2A). Over time, there was progressive decline in alpha-diversity after adjusting for sample type (rectal swab vs. stool samples) (Figures 2A and S2).

**Figure 2.**
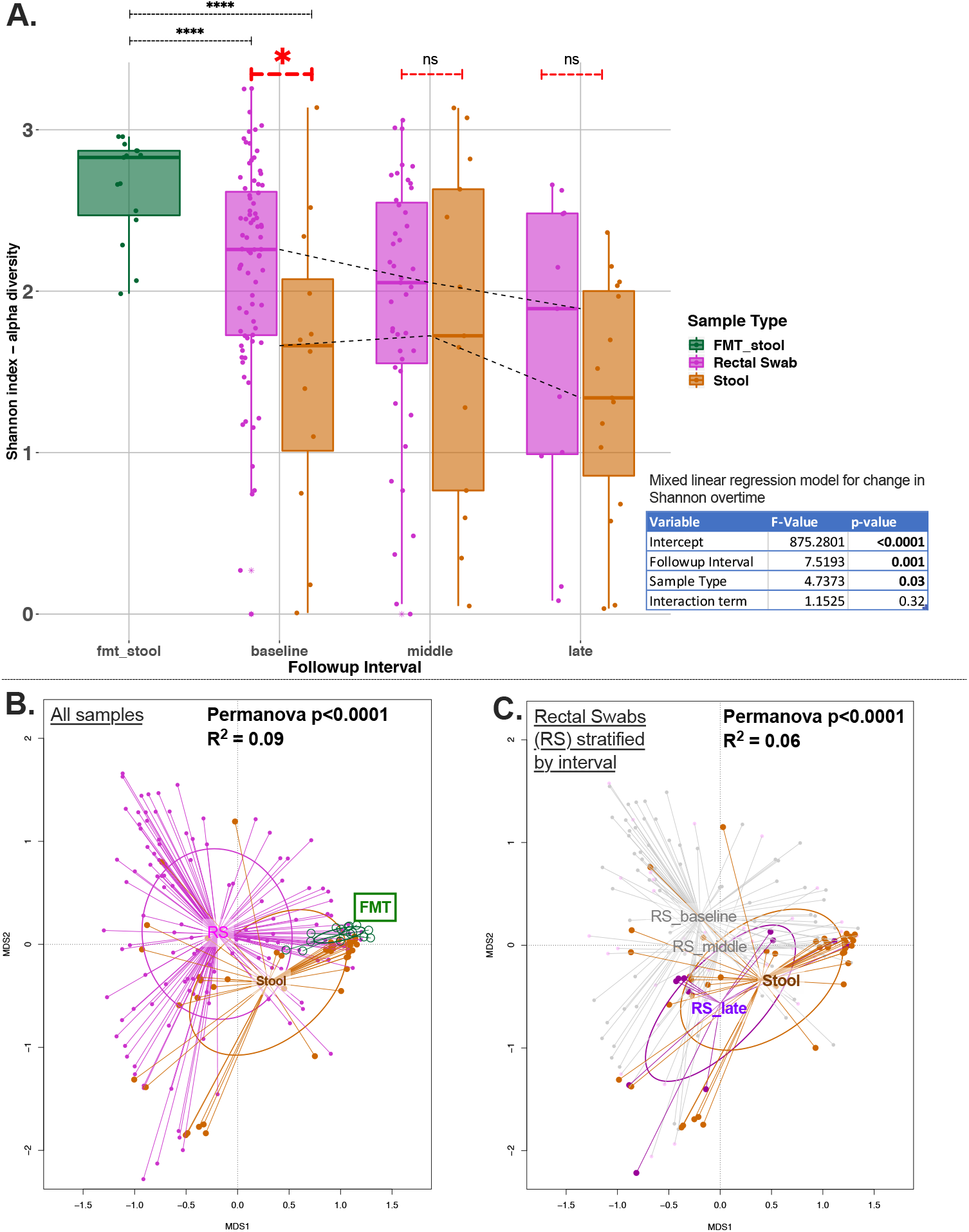
Alpha and beta-diversity comparisons show markedly different representations of the gut microbiome by sample type. Panel A – Alpha diversity analyses by sample type and follow up interval showed that rectal swabs had higher Shannon index than stool samples at the baseline timepoint by a Wilcoxon test (p<0.02) and not in subsequent follow up intervals. Both rectal swabs and stool samples at baseline had significantly lower alpha diversity than FMT samples (p<0.0001). There was significant decline of Shannon index over time, adjusting for sample type with a mixed linear regression model with random patient intercepts (shown in Table). Panel B – Beta Diversity analyses: Principal coordinates analyses of Bray-Curtis dissimilarity indices between rectal swabs and stool samples. Greater distance between samples indicates greater dissimilarity. In the left panel, all available samples are stratified by sample type, showing significant differences between rectal swabs and stool samples (permutation analysis of variance p=0.0001). FMT samples appeared compositionally more similar to stool samples rather than rectal swabs from critically-ill patients. In the right panel, stratified analyses by study follow-up interval for rectal swabs shows that rectal swabs in the late interval are more similar to stool samples (overlapping ellipsoids) compared to rectal swabs obtained earlier (baseline or middle interval).

Rectal swabs and stool samples were systematically different by beta-diversity (Bray-Curtis dissimilarity index [Permutational Multivariate Analysis of Variance-PERMANOVA] p<0.0001, Figure 2B), even after adjustment for potential confounders (Table S2). Of note, FMT stool samples were more similar by beta-diversity to stool samples from ICU patients than rectal swabs (Figure 2B). By stratifying rectal swabs based on follow-up interval, visualization of beta-diversity with Principal Coordinates Analyses (PCoA) revealed that rectal swabs in the late interval were compositionally more similar to stool samples than rectal swabs obtained earlier (Figure 2C). By PERMANOVA, a statistically significant temporal effect for changes on beta-diversity was found only for rectal swabs (p=0.002) and not for stool samples (Table S3). Next, in the subset of patients with both stool and rectal swabs available at different intervals, we examined the relative impact of the patient identity vs. the sample type variable on beta-diversity (effectively asking the question whether different sample types obtained from the same patients were more similar to each other compared to same sample types obtained from different patients). By PERMANOVA, the sample type was the only variable significantly associated with differences in beta-diversity (p=0.002) (Table S4), i.e. knowing whether a community taxonomic profile was derived from a rectal swab vs. a stool sample was more important than knowing from which patient this sample was taken from. In further sensitivity analyses, we examined for the potential impact of different methods of stool sample acquisition (collected from a rectal tube bag vs. from bowel movements) but did not find any significant alpha or beta-diversity differences (Figure S3).

Analyzing the taxonomic composition at the phyla level, we noted that rectal swabs had higher relative abundance of Actinobacteria (a commensal skin microbe) compared to stool samples (Wilcoxon p=0.05 for additive log ratio transformation comparison), but the Actinobacteria abundance declined significantly over time (Figure S4). At the genus level, stool samples had higher relative abundance of *Akkermansia, Bacteroides, Enterococcus* and *Parabacteroides* taxa, which are considered typical members of the gut microbiome in critically-ill patients (Figure S5).

Examination of the taxonomic composition at the genus level for 10 patients who had both sample types available at different follow-up intervals showed marked discordance between rectal swabs vs. stool samples (Figure S6).

## Discussion

Our analyses in a large cohort of critically-ill patients highlight significant differences between the more accessible rectal swabs and the harder to obtain, but commonly viewed as reference-standard stool samples. In our study, stool sample availability captured distinct patient characteristics, perhaps because morbid critically-ill patients with longer ICU stays had higher likelihood of stool passage and collection during the study follow-up period. Nevertheless, stool samples and rectal swabs had significant differences in alpha and beta-diversity even after adjustment for potential confounders of the associations between sample types and microbiota community composition.

Systematic differences in alpha and beta-diversity by sample type were largely attributable to the baseline samples acquired close to ICU admission. At this early timepoint, 87% of the available samples were rectal swabs and most patients were not receiving enteral nutrition (58%). Stool presence in the rectal vault may have been limited, leading to “unsoiled” swabbing of the rectal mucosa and peri-rectal skin, with a resulting disproportionate abundance of skin bacteria (i.e. Actinobacteria phyla) than what would be expected for gut microbiota profiles (11). The temporal convergence of microbial profiles between rectal swabs and stool samples observed in our study suggests that progressive recovery of gut motility during the ICU course and stool presence in the rectal vault may have improved the reliability of “soiled” rectal swab sampling, although we did not qualitatively score the macroscopic appearance of swabs as in the study by Bansal et al. (9). In addition, our study design with periodic sampling in predefined intervals rather than on consecutive days has hindered our ability to detect day-to-day, dynamic changes of gut microbiota communities and limited the number of follow-up samples in our cohort. Thus, analyses in the late interval have low statistical power and may have also been affected by informative censoring, i.e. patients not contributing late follow-up samples due to rapid clinical improvement and discharge from the ICU or due to clinical deterioration and early ICU death. Since we were not able to perform head-to-head comparisons of rectal swabs and stool samples obtained at the same time, the notable discordance of microbial community profiles by sample type observed in the small subset of patients with longitudinal samples of both types requires cautious interpretation, given that it was not possible to account for patient-level temporal variability.

Our results call for caution in the design of gut microbiome studies in critically-ill patients. Despite their availability, rectal swabs may offer biased representations of the presumed gut microbial communities, especially when conducted early in the course of critical illness. Consequently, analyses of gut microbiota studies need to take into account both the actual sample types used and the timing of sample acquisition, because rectal swabs and stool samples are not interchangeable for the purpose of microbiota sequencing profiling. Accurate and reproducible delineation of the role of the gut microbiome in critical illness will require consistent sampling protocols, clinical variable recording and longitudinal assessments.

## Data Availability Statement

Raw sequences used for this project have been deposited and are publicly available at http://www.ncbi.nlm.nih.gov/bioproject/516701 Taxa tables, metadata and R statistical code required for the conduct of the analyses described here in have been deposited and are publicly available at https://github.com/MicrobiomeALIR/

