## Supplemental Methods, Tables, and Figures for "Rectal swabs in critically-ill patients provide discordant representations of the gut microbiome compared to stool samples: a brief methodologic report"

**Supplement**

**Contents:**


**Expanded Methods:** pages 2-8

**Results:** pages 9-22

**References:** pages 23-24

**Expanded Methods**

Clinical cohort:

From March 2015 – August 2017, we prospectively enrolled consecutive mechanically-ventilated patients with acute respiratory failure from the Medical Intensive Care Unit (ICU) at the University of Pittsburgh Medical Center (UPMC) into the Pittsburgh Acute Lung Injury Registry and Biospecimen Repository (ALIR). Eligible patients were 18 years or older with acute respiratory failure who were intubated and mechanically-ventilated. Exclusion criteria included inability to obtain informed consent, presence of tracheostomy, or mechanical ventilation for more than 72 hours prior to enrollment. The ALIR study was approved by the University of Pittsburgh Institutional Review Board (protocol PRO10110387), and written informed consent was provided by all participants or their surrogates in accordance with the Declaration of Helsinki.

For comparisons with healthy controls, we also included samples from donated stool for fecal microbiota transplant from 15 healthy adult donors.

**Sample Collection:**

From consented critically-ill patients, we collected rectal swabs and/or stool samples at three time intervals starting at the time of intubation and continuing up to 10 days if the patient remained in the ICU (defined as baseline [days 0-2], middle [days 3-6] and late intervals of follow-up [days 7-10]). Rectal swabs were collected as per routine clinical practices (i.e. placing the patient in a lateral position, entering the tip of the 6" cotton tip swab in the rectal canal and rotating gently for 5 seconds), unless clinical reasons precluded movement of the patient (e.g. severe hemodynamic or respiratory instability). Stool samples were collected when available, either by taking a small sample from an expelled bowel movement (before cleaning of the patient and disposal of stool), or through a fecal management system (rectal tube) placed for management of diarrhea and liquid stool collection.

Stool samples or rectal swabs were placed in sterile collection tubes and then capped, labeled and stored at -80°C until further processing.

**Clinical Data Recording:**

Detailed data was obtained from the electronic medical record on the day of enrollment including:

1) Demographic information: age, sex, height, weight, body mass index
2) Laboratory results: white blood cell count, hemoglobin, platelets, serum bicarbonate, arterial pH, partial pressure of oxygen, partial pressure of carbon dioxide.
3) Mechanical ventilation parameters: respiratory rate, tidal volume (normalized per kilogram ideal body weight), positive end-expiratory pressure, peak inspiratory pressure, and fraction inspired oxygen concentration.
4) Severity of illness scores: sequential organ failure assessment (SOFA) score excluding the neurologic component because all patients were intubated and receiving sedative medications, impairing our ability to perform assessment of the Glasgow Coma Scale in a consistent and reproducible fashion) (1). SOFA score was calculated by using the physiologically worse values within 24hrs of enrollment.
5) Lung Injury Prediction Scores (2).
6) Medication administration: antibiotics, vasopressors, and sedatives administered during the first week of ICU course from intubation.

7) Gastrointestinal/Nutritional Factors: enteral nutrition administration, number of bowel movements or clinical diarrhea (as assessed by bedside nurses), and rectal tube presence.
8) Outcomes: length of ICU stay, 30-day mortality, incident shock, sepsis, and occurrence of acute kidney injury (AKI) within the first week of ICU stay(4).

### Laboratory Methods:

#### DNA extraction:

We extracted bacterial DNA directly from samples (rectal swabs and stool samples) using Powersoil (Qiagen) extraction kit following manufacturer's instructions, as previously described (3). For rectal swabs, we agitated the cotton tip in the extraction column tube for e1-2 min and until visible particles of the of biospecimen were released from the cotton tip into the C1 solution of the Powersoil kit. For stool samples, we used 30-40mg of stool sample directly inoculated into the C1 solution of the extraction column tube. As DNA extraction negative controls, we added DNA-free sterile water in one extraction column per each batch of clinical sample DNA extractions (~12-15 samples per batch).

#### 16S rRNA gene sequencing:

We amplified extracted DNA by PCR using the method of Caporaso et al. and the Q5 HS High-Fidelity polymerase (NEB) targeting the V4 hypervariable region of the 16S rRNA gene (5). We utilized reagent controls for each step of the process (DNA extraction and PCR amplification as described above). We amplified four microliters per reaction of each sample with a single barcode in triplicate 25 microliter reactions. As PCR-negative controls, we added DNA-free sterile water in the PCR reaction mix in amount equal to the amount of template DNA from clinical samples used (typically 6 microliters). As PCR amplification positive controls, we utilized the ZymoBIOMICs Microbial Community DNA Standard (Zymo Research, Irvine, CA), a mock microbial community consisting of genomic DNA of eight bacterial strains. We utilized 1 microliter of the genomic mixture with concentration of 10ng/microliter for each reaction. Cycle conditions were 98˚C for 30s, then 33 cycles of 98˚C for 10s, 57˚C for 30s, 72˚C for 30s, with a final extension step of 72˚C for 2 min. We combined triplicates and purified with the AMPure XP beads (Beckman) at a 0.8:1 ratio (beads:DNA) to remove primer-dimers. We quantitated eluted DNA on a Qubit fluorimeter (Life Technologies). We performed sample pooling on ice by combining 20 ng of each purified band. For negative controls and poorly performing samples, we used 20 microliters of each sample. We purified the sample pool with the MinElute PCR purification kit. The final sample pool underwent two more purifications – AMPure XP beads to 0.8:1 to remove all traces of primer dimers and a final cleanup in Purelink PCR Purification Kit (Life Technologies). We quantitated the purified pool in triplicate on the Qubit fluorimeter prior to preparing for sequencing. We prepared the sequencing pool according to instructions by Illumina, with an added incubation at 95˚C for 2 minutes immediately following the initial dilution to 20 picomolar. We then diluted the sequencing pool to a final concentration of 7 pM + 15% PhiX control. Amplicons were sequenced on the Miseq platform.

**Analytics**:

16S Sequence quality control:

Sequences from the pooled sequencing run were demultiplexed into individual sample/replicate fastq files. Each fastq file was then processed through the Center for Medicine and the Microbiome (CMM) custom modular read QC pipeline that was configured to perform the following steps: low complexity filtering, QV trimming, Illumina sequencing adapter trimming, and 16S primer trimming. Low complexity filtering utilized NCBI BLAST's dustmasker (6). Reads with greater than 80% low complexity regions were filtered. QV trimming and filtering utilized the FASTX Toolkit. The trimming threshold was QV>25 from the 3' end, and length>125 bp after trimming. The subsequent filtering threshold was >25 across >95% of the read. The Illumina sequencing adapter and 16S primer trimming was performed with cutadapt (7).

<u>16S Clustering and annotation</u>**:**

Paired sequences with forward and reverse reads passing the QC filtering and trimming steps were then mated (end aligned and a consensus sequence computed) using the make.contigs function of Mothur. Consensus sequences were screened to limit the overlap mismatch to no more than 20%. The maximum number of N's allowed in the overlap was 4 and the minimum overlap was required to be greater than 25bp. Consensus sequences passing screening were then passed through CMM's 16S clustering and annotation pipeline, a Mothur-dependent wrapper designed to streamline and automate the execution of the following Mothur steps: unique.seqs, align.seqs, screen.seqs, filter.seqs, second uniq.seqs, pre.cluster, chimera.uchime, remove.seqs, classify.seqs, dist.seqs, cluster, make.shared, and classify.otu. Mothur output files were then reformatted to sample x category (taxonomic levels or OTU) matrices for downstream statistical modeling (8).

<u>Taxa table edits</u>:

The taxa table was filtered for low abundance taxa (relative abundance, <0.005%) and singletons. We did not filter clinical samples for any taxa detected in the negative control samples. Five samples were subjected to repeat PCR amplification due to poor performance in initial reactions, and were then removed from analyses as duplicates.

Statistical Analysis

We calculated descriptive statistics of baseline characteristics and performed non-parametric comparisons (Wilcoxon test for continuous and Fisher's exact test for categorical variables) using the R software (R Foundation for Statistical Computing, 2016). We performed ecological analyses of alpha-diversity (richness-Shannon), beta diversity (Bray–Curtis dissimilarity index), and taxonomic relative abundance at the phyla and genus level with R *vegan* package (9). Beta-diversity comparisons with permutation analysis of variance (Permanova at 1000 permutations) were visualized with Principal Coordinates Analyses. Taxonomic abundance differences between groups were calculated following transformation of abundance data with the additive log ratio, performed for the top 4 phyla (Firmicutes, Bacteroidetes, Actinobacteria and Proteobacteria) and the top 20 genera (10).

In order to assess for longitudinal changes of alpha-diversity over time as well as to account for the effects of potential confounders on the associations between sample type and gut microbiota profiles, we constructed a set of multivariate models as described below.

As confounders, we considered potential variables that may impact gut motility and bowel content in critically-ill patients, as well as having potential associations with the gut microbiome:

1. Early Enteral Nutrition (defined as initiation of gastric or enteral feeds within 48hrs of ICU admission).
2. SOFA score on continuous scale, as a marker of severity of illness.
3. Presence of rectal tube, as an indicator of severe diarrhea requiring a fecal management system.

4. Obesity, defined as a BMI>30, which has been associated with gut microbiome profiles, but for practical purposes may also affect our ability to perform rectal swabbing especially in the subset of patients with morbid obesity.
5. Clinical diagnosis of sepsis, associated with administration of broad-spectrum antibiotics, which can cause diarrhea, but may also affect the gut microbial communities.

We then constructed the following multivariate models:

1. Alpha-diversity in baseline samples: we constructed a mixed linear regression model with Shannon index as the response variable, random patient intercepts, and then the sample type as an independent variable adjusted for these 5 confounders.
2. Alpha-diversity over time: we constructed mixed linear regression models considering random patient-level intercepts and examined for the independent effects of ordered follow-up intervals (baseline vs. middle vs. late) after adjusting for sample type (rectal swab vs. stool samples) for all patients as well as for the subset of patients with available follow-up samples (i.e. excluding patients with baseline samples only).
3. Beta-diversity: we performed PERMANOVA analyses with beta-diversity as the dependent variable and the sample type as an independent variable adjusted for the 5 confounding variables described above.

### Supplemental Results

**Figure S1: Number of reads (high quality 16S rRNA gene sequences) by sample type.** Clinical samples are shown with colored filled boxplots (green for stool samples from healthy donors of fecal microbiota transplant specimens, pink for rectal swabs from critically-ill patients and brown for stool samples from critically-ill patients) whereas experimental controls are shown with empty boxplots. Median and interquartile range (IQR) of number of reads for each sample type are shown with blue fonts. Comparisons of number of reads between samples were performed with nonparametric Wilcoxon tests. Gut microbiome specimens from critically-ill patients (rectal swabs or stool samples) had numbers of reads in the range of polymerase chain reaction (PCR) positive controls (pcrpos), but much higher numbers of reads compared to negative controls (negative controls for DNA extractions (mobioextrneg) and PCR negative controls (pcrneg), as well as compared to fecal microbiota transplant speciments (fmt_stool). There was no difference in median number of reads between rectal swabs (median 4270, IQR 1022) and stool samples (median 4185, IQR 1079, p=0.25).

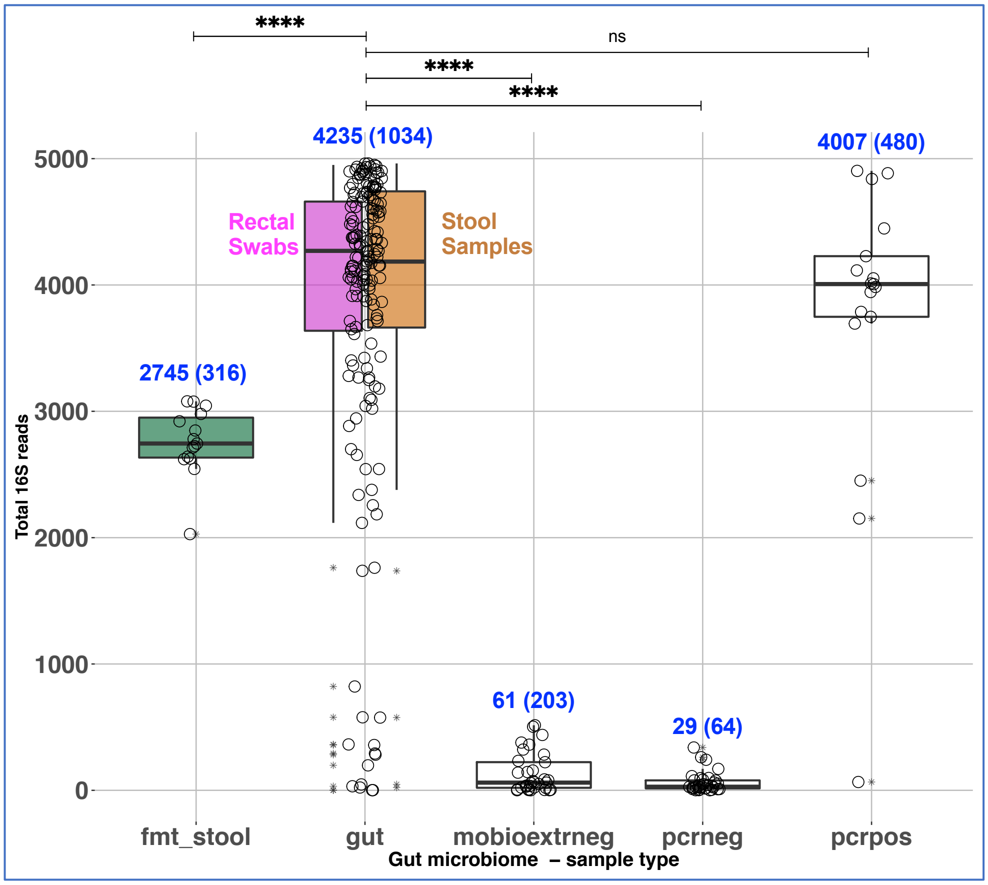

****
ns
****
****
4235 (1034)
4007 (480)
2745 (316)
Rectal Swabs
Stool Samples
61 (203)
29 (64)
5000
4000
3000
2000
1000
0
Total 16S reads
fmt_stool
gut
mobioextrneg
pcrneg
pcrpos
Gut microbiome – sample type

**Table S1: Results of mixed linear regression model for independent effects of sample type on baseline sample alpha-diversity (Shannon Index).** Statistically significant associations ($p<0.05$) are shown in bold. Sample type was significantly associated with alpha-diversity after adjusting for potential confounders.

| Variable | F-value | p-value |
|---|---|---|
| **Sample type (rectal swab vs. stool sample)** | 8.88 | **0.004** |
| **SOFA score** | 0.697 | 0.41 |
| **Clinical Sepsis** | 0.309 | 0.58 |
| **Presence of rectal tube** | 6.24 | **0.003** |
| **Early enteral nutrition** | 2.09 | 0.15 |
| **Obesity** | 0.989 | 0.32 |

**Figure S2. Significant decline in alpha diversity (Shannon index) over time independent of sample type for the subset of patients (n=27) with available samples at all follow-up intervals.** No significant differences in Shannon index between rectal swabs and stool samples at each interval. There was significant decline of Shannon index overtime, adjusting for sample type with a mixed linear regression model with random patient intercepts (shown in Table).

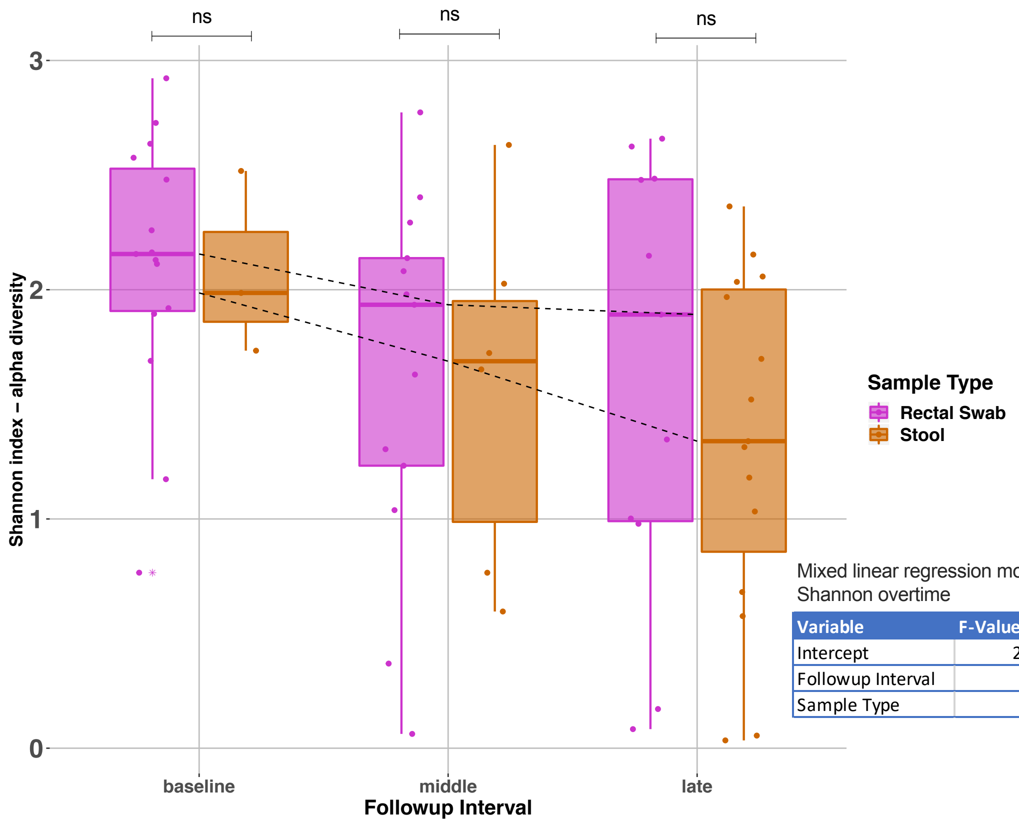


Mixed linear regression model for change in Shannon overtime

| Variable | F-Value | p-value |
|---|---|---|
| Intercept | 207.2 | **<0.0001** |
| Followup Interval | 6.1 | **0.005** |
| Sample Type | 0.3 | 0.55 |

**Table S2: Significant independent associations of sample type with beta-diversity differences based on Permutational Multivariate Analysis of Variance models accounting for effects of potential confounders**. Table S2A includes results of the model considering all samples available and table S2B considering only baseline samples. Statistically significant associations ($p<0.05$) are highlighted in bold.

| A: all samples available | $R^2$ | p-value |
|---|---|---|
| Sample type (rectal swab vs. stool sample) | 0.04 | **0.001** |
| Early enteral nutrition | 0.005 | 0.51 |
| SOFA Score | 0.005 | 0.58 |
| Presence of rectal tube | 0.021 | **0.006** |
| Obesity | 0.007 | 0.12 |
| Clinical sepsis | 0.008 | 0.09 |

| B: baseline samples only | $R^2$ | p-value |
|---|---|---|
| Sample type (rectal swab vs. stool sample) | 0.021 | **0.023** |
| Early enteral nutrition | 0.018 | **0.039** |
| SOFA Score | 0.009 | 0.67 |
| Presence of rectal tube | 0.04 | **0.004** |
| Obesity | 0.011 | 0.34 |
| Clinical sepsis | 0.008 | 0.09 |

**Table S3: Permutational Multivariate Analysis of Variance models for assessment of temporal changes of beta-diversity separately in rectal swab samples (Table S3A) and stool samples (Table S3B).** Significant changes over time were observed only in the rectal swab subgroup.

| A: Rectal Swabs | $R^2$ | p-value |
|---|---|---|
| Follow-up interval | 0.031 | **0.002** |
| **B: Stool samples** | **$R^2$** | **p-value** |
| Follow-up interval | 0.05 | 0.40 |

**Table S4: Permutational Multivariate Analysis of Variance model for assessment of the impact of patient identity vs. sample type on beta-diversity.** The sample type variable was the only one significantly associated with beta-diversity differences (p<0.0001), i.e. knowing whether a community taxonomic profile emerged from a rectal swab vs. a stool sample was more important than knowing from which patient this sample was taken from.

| Variables | $R^2$ | p-value |
|---|---|---|
| **Sample type** | 0.09 | **0.002** |
| **Subject ID** | 0.03 | 0.49 |

**Figure S3. No significant alpha (A) or beta-diversity (B) differences between stool samples obtained from patient bowel movements vs. liquid stool collected from rectal tube bags**.

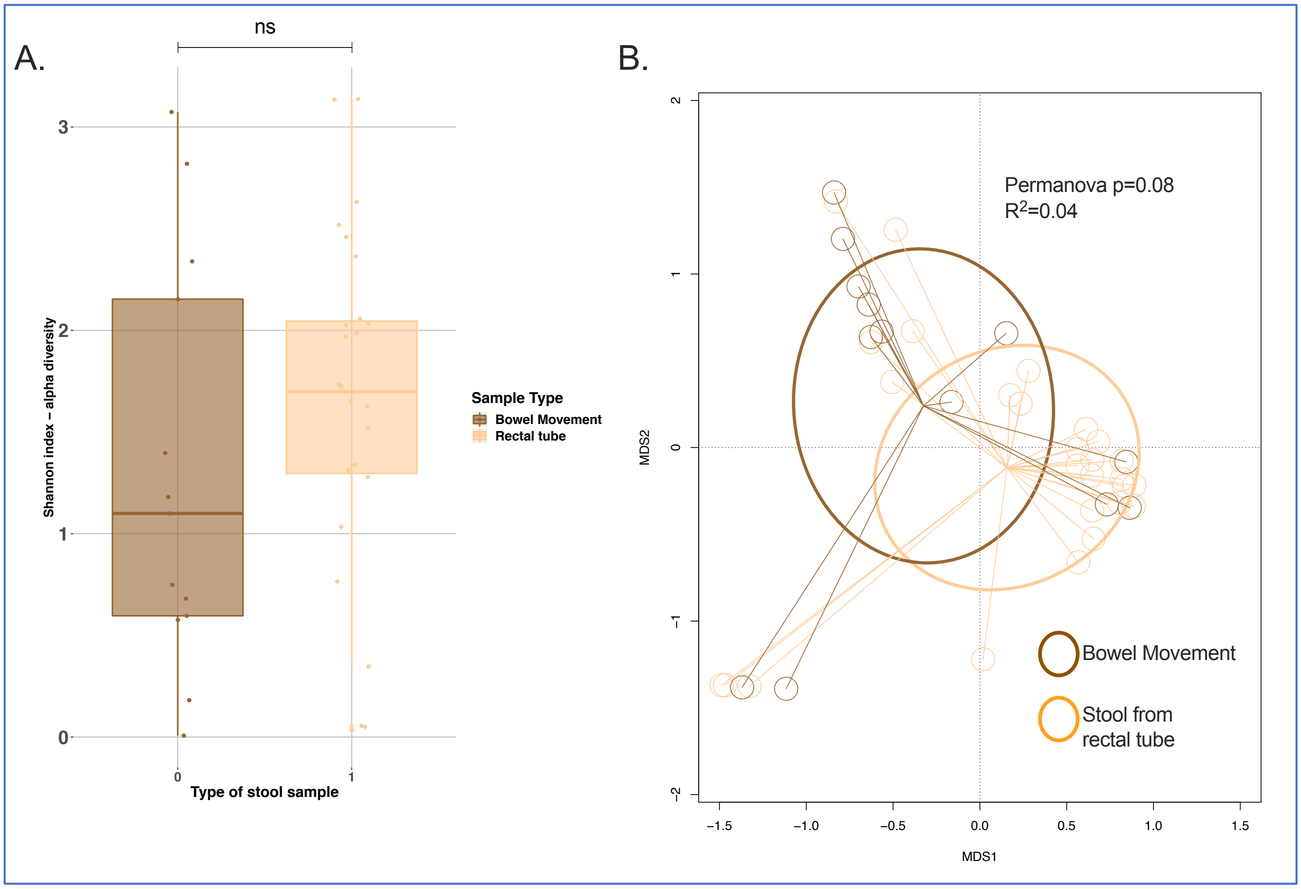

**Figure S4.** Taxonomic composition at the phyla level for rectal swabs, stool samples and FMT samples (Panel A). Relative abundance for the 4 most common phyla (Firmicutes, Bacteroidetes, Proteobacteria and Actinobacteria) is shown, broken down by follow-up interval for rectal swabs and stool samples. Additive log-ratio transformed abundance of Actinobacteria overtime in rectal swabs (Panel B) and stool samples (Panel C). Significant decline in Actinobacteria abundance was evident for the rectal swabs only, examined by mixed linear regression model with study follow-up interval as an independent variable and additive log ratio of abundance as the response variable, with random patient intercepts.

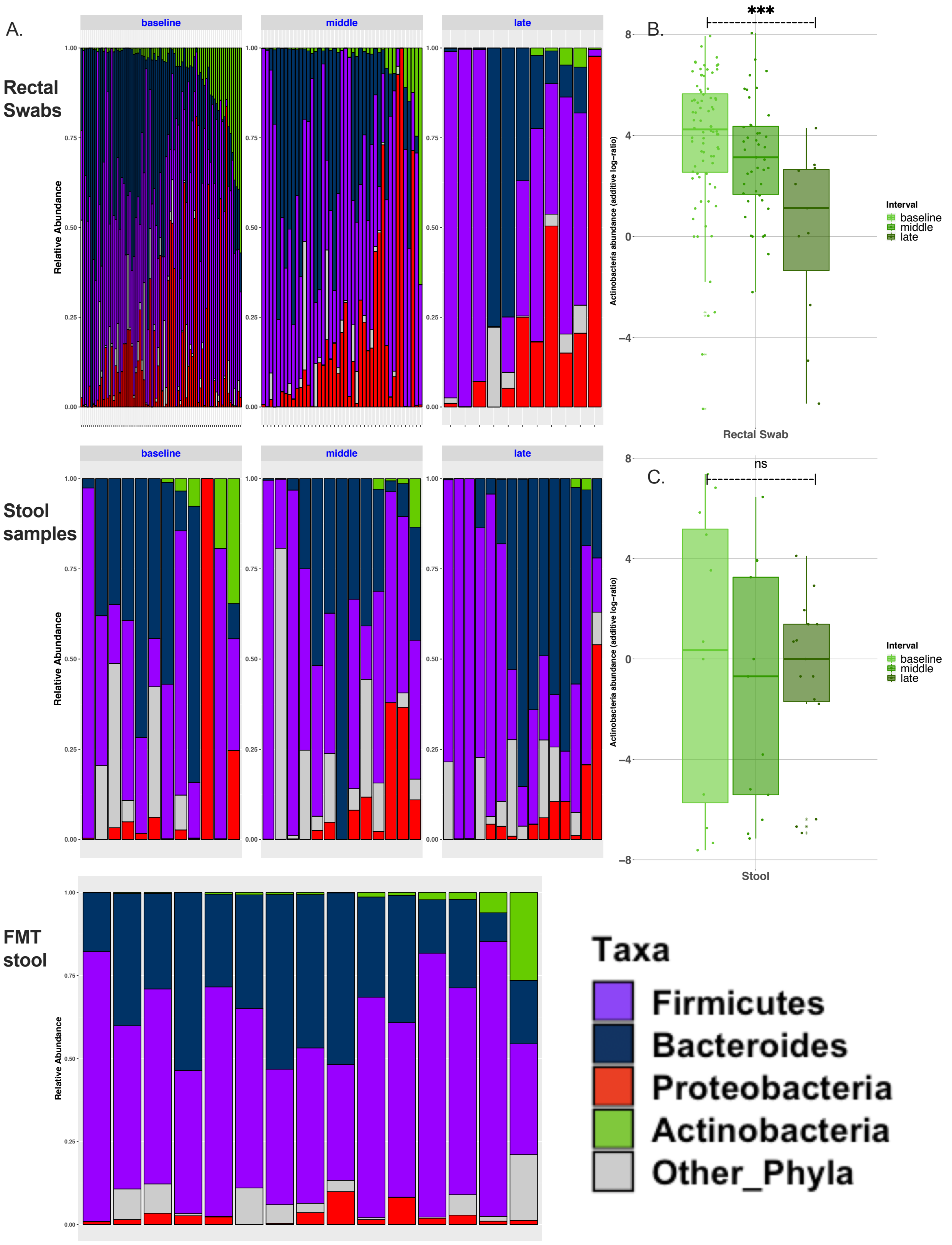
A.
Rectal Swabs
Stool samples
FMT stool
baseline
middle
late
Relative Abundance
B.
C.
***
ns
Actinobacteria abundance (additive log−ratio)
Interval
baseline
middle
late
Rectal Swab
Stool
Taxa
Firmicutes
Bacteroides
Proteobacteria
Actinobacteria
Other_Phyla

**Figure S5**: Taxa relative abundance box plot by sample type for 15 most common taxa and other genera. Significantly increased relative abundances in stool vs. rectal swabs are displayed in the *Akkermansia*, *Bacteroides*, *Enterococcus*, and *Parabacteroides* taxa. Comparisons were done with Wilcoxon tests on additive log-ratio transformed abundances of individual genera. Statistical significance is shown with asterisks (ns: not significant; *: $p<0.05$; **: $p<0.01$; ***: $p<0.001$; ****: $p<0.0001$).

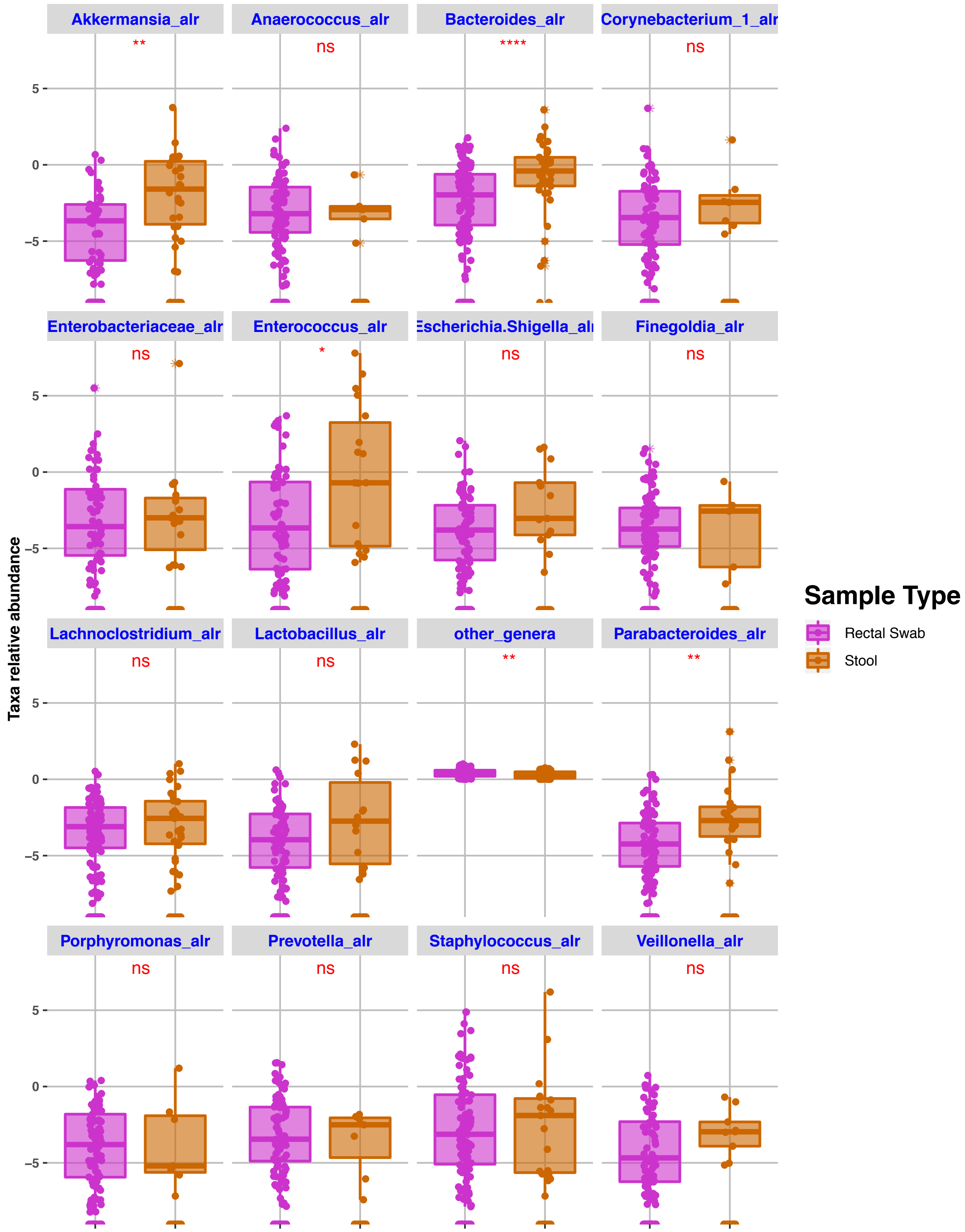
Akkermansia_alr
**
Anaerococcus_alr
ns
Bacteroides_alr
****
Corynebacterium_1_alr
ns
Enterobacteriaceae_alr
ns
Enterococcus_alr
*
Escherichia.Shigella_alr
ns
Finegoldia_alr
ns
Lachnoclostridium_alr
ns
Lactobacillus_alr
ns
other_genera
**
Parabacteroides_alr
**
Porphyromonas_alr
ns
Prevotella_alr
ns
Staphylococcus_alr
ns
Veillonella_alr
ns
Taxa relative abundance
5
0
−5
Sample Type
Rectal Swab
Stool

**Figure S6**: **Taxonomic composition bar plot at the genus level for 10 patients with both stool samples and rectal swabs available at different time points.** Rectal swabs are shown in the top row and stool samples in the bottom row, with each patient unique 4-digit ID highlighting samples belonging to the same patient. The relative abundances for the 15 most common taxa are shown with different colors, and the added relative abundance of all other component taxa is shown in gray ("other genera"). Marked discordance between rectal swabs and stool sample taxonomic composition is demonstrated.

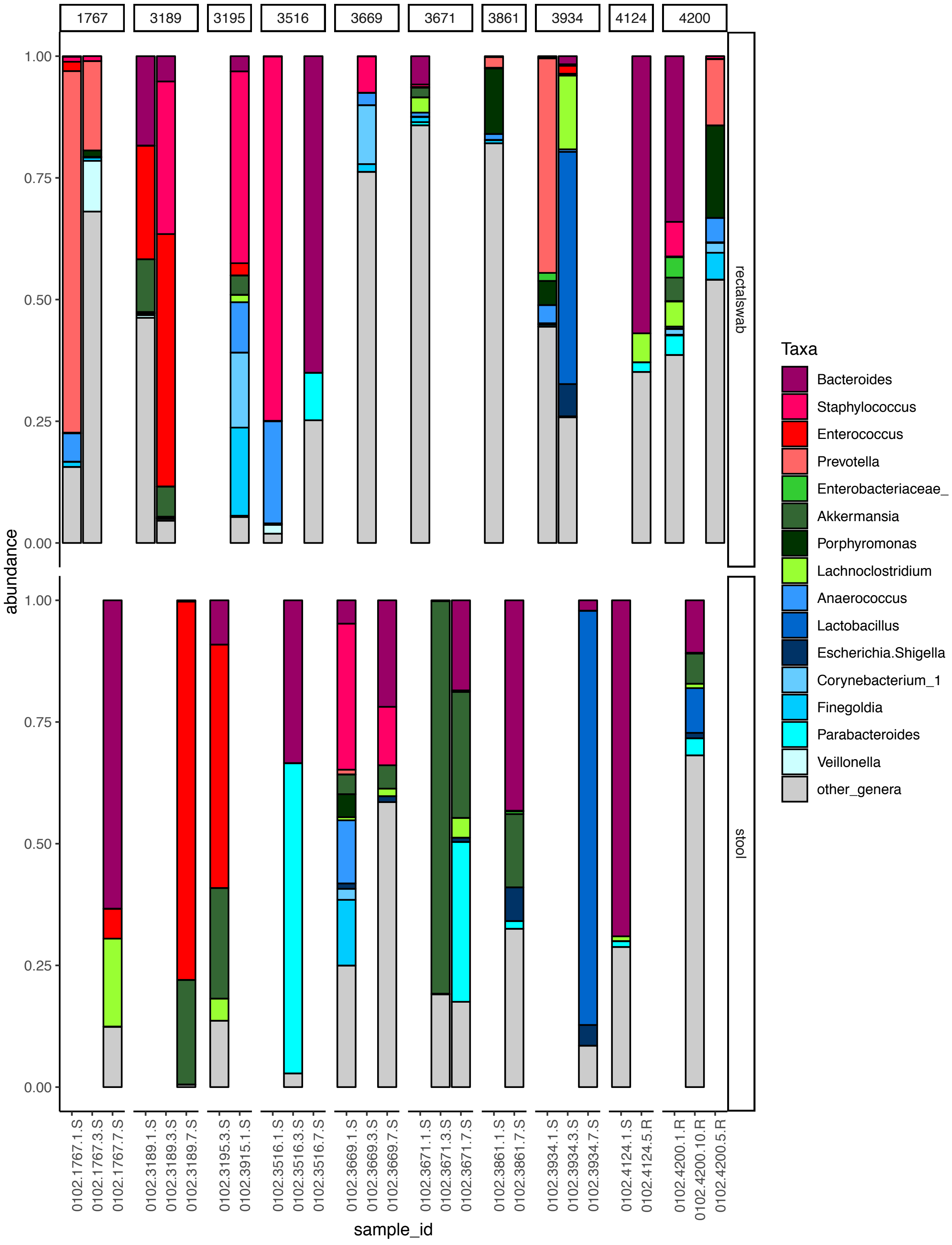

1767
3189
3195
3516
3669
3671
3861
3934
4124
4200
abundance
1.00
0.75
0.50
0.25
0.00
rectalswab
stool
0102.1767.1.S
0102.1767.3.S
0102.1767.7.S
0102.3189.1.S
0102.3189.3.S
0102.3189.7.S
0102.3195.3.S
0102.3915.1.S
0102.3516.1.S
0102.3516.3.S
0102.3516.7.S
0102.3669.1.S
0102.3669.3.S
0102.3669.7.S
0102.3671.1.S
0102.3671.3.S
0102.3671.7.S
0102.3861.1.S
0102.3861.7.S
0102.3934.1.S
0102.3934.3.S
0102.3934.7.S
0102.4124.1.S
0102.4124.5.R
0102.4200.1.R
0102.4200.10.R
0102.4200.5.R
sample_id
Taxa
Bacteroides
Staphylococcus
Enterococcus
Prevotella
Enterobacteriaceae_
Akkermansia
Porphyromonas
Lachnoclostridium
Anaerococcus
Lactobacillus
Escherichia.Shigella
Corynebacterium_1
Finegoldia
Parabacteroides
Veillonella
other_genera
